# Quantification of single cell-type-specific alternative transcript initiation

**DOI:** 10.1101/2025.04.29.651292

**Authors:** Shuhua Fu, Parker Wilson, Bo Zhang

## Abstract

Cells can transcribe different isoforms of a gene by using distinct Transcriptional Start Regions (TSRs), which are recognized by RNA-Polymerase II and regulated by cell-type-specific expressed transcription factors, eventually forming tissue and cell-type-specific expression during development. However, how the distinct TSRs are selectively activated in different tissues and cell types remains largely uncharacterized. To better explore the alternative usage of gene TSRs, we developed *TSRdetector*, a novel bioinformatic method specifically designed to detect the significant usage alteration of gene TSRs in different tissues, cell types, or diseases. *TSRdetector* can process either scRNA-seq or bulk-RNA-seq transcriptome data, define dominant TSRs, and compute the differential usage of gene TSRs between given conditions. To demonstrate the capacity of *TSRdetector*, we applied *TSRdetector* to analyze a 10X snRNA-seq dataset of healthy and diabetic human kidneys, a Smart-seq2 dataset of mouse B-cell differentiation, and bulk-RNA-seq data of pluripotency transition in human ESCs. In all three analyses, TSRdetectors discovered significant alterations in TSR usage, accompanied by the significant remodeling of the epigenetic landscape. Alteration of TSR usage can change the dominant transcript isoform and further affect the major protein products. Interestingly, a large proportion of these alterations of TSR usage in different cell types did not change the overall gene expression, revealing unique transcription regulations that are independent of expression level. In summary, *TSRdetector* is a user-friendly package to analyze the differential usage of gene TSRs by using both scRNA-seq and bulk-RNA-seq data, and can be used to explore the alternative transcription initiations of genes at the single cell-type level.

## INTRODUCTION

The complex gene structure in eukaryotic genomes allows for multi-level regulation, including alternative splicing and alternative transcription start region (TSR) usage, producing transcript isoforms with distinct lengths and structures(Singer et al. 2008; Zavolan et al. 2003). Alternative TSR usage is widespread but highly dynamic across tissues, cell types, and healthy states (Alasoo et al. 2019). *Sst2* can utilize distinct TSRs in different tissues: TSR in exon 1 is active in AtT-20 tumor cells, exon 2 TSR in brain and pancreas, and exon 3 TSR in lung and kidney(Kraus et al. 1998). Novel alternative first exon usages in humans and mice were also found during inflammation using short-read and Nanopore long-read sequencing(Robinson et al. 2021; Ueda et al. 2024). In human skeletal muscle, 10% of genes changed expression post-stress due to alternative TSR activation(Makhnovskii et al. 2022). Alternative TSR usage also plays a role in environmental and stress responses. In Arabidopsis, TSR shifts in *GLYK* regulate protein localization under different lighting conditions(Ushijima et al. 2017). In mouse striatum, chronic cocaine exposure alters TSR usage to increase gene expression(Mains et al. 2011). The *Frq* gene, a core component of the circadian clock in Neurospora crassa, switches TSRs in response to light and temperature changes (Colot et al. 2005). Recently, 3’ end site selection was found to couple with 5’ alternative TSR usage in Drosophila and the human nervous system, linking distinct mRNA isoform expression to TSR selection(Alfonso-Gonzalez et al. 2023). Transposable elements (TEs) also contribute to TSR regulation. An MT2B2 retrotransposon-derived promoter generates an N-terminally truncated *Cdk2ap1* in preimplantation embryos, affecting cell proliferation(Modzelewski et al. 2021). 453 TEs were identified as TSRs to initiate the host gene transcription during mouse embryonic development(Miao et al. 2020), while human airway epithelial cells use a TE-derived TSR to transcribe *IL33* in chronic airway disease(Raphael et al. 2024). In a recent TCGA pan-cancer analysis, 1,068 TE-derived TSRs were identified to generate tumor-specific TE-chimeric antigens(Shah et al. 2023). Individual-specific TSR alterations were further linked to patient survival in TCGA data (Demircioglu et al. 2019). These findings underscore alternative TSR usage as a crucial regulatory mechanism across species, influencing development, disease, and environmental adaptation.

RNA sequencing, a widely used method, provides comprehensive insights into complex RNA transcripts and is crucial for studying alternative splicing and TSR usage in a high-throughput manner. Different tools have been developed to analyze alternative splicing and TSRs/promoter usage, such as Cufflinks(Trapnell et al. 2010) and JuncBASE(Brooks et al. 2011), can identify RNA isoforms, allowing TSR definition after transcript assembly. SEASTAR was developed to quantify first exon usage via de novo assembly of bulk RNA-seq data, revealing significant TSR shifts during fibroblast reprogramming into iPSCs(Qin et al. 2018). TappAS is a framework for identifying splicing patterns and alternative TSRs using short- and long-read sequencing(de la Fuente et al. 2020). Cass *et al*. developed mountainClimber to annotate alternative TSRs and polyadenylation sites in parallel(Cass and Xiao 2019). Recent advancements in experimental techniques have enabled single-cell and single-nucleus RNA sequencing, providing detailed transcriptome insights at the single-cell level. These methods generate vast datasets capturing complex RNA transcripts. Karlsson et al. observed a high correlation in alternative TSR expression patterns in single cells of the same mouse brain cell type (Karlsson et al. 2017). CamoTSS, a recently developed tool, creatively leveraged the 5’ tag-based scRNA-seq data to identify TSSs and quantify their expression(Hou et al. 2023).

In this article, we introduce TSRdetector, a tool designed for single-cell transcriptome data, to detect and quantify alternative transcription start region (TSR) usage. TSRdetector can first leverage the long-read RNA-seq sequencing dataset to provide high-quality transcript annotation, and further quantify the expression of transcripts in single-cell transcriptome data, calculate the TSR usage based on overall gene expression at single-cell type resolution. TSRdetector can be used to process the RNA-seq data from platforms such as 10x Genomics Chromium (5’ protocol, SMRT-seq), Smart-Seq2/3, and other sequencing methods that enriched 5’ end transcript information and gene-body splicing information, including high-quality bulk RNA-seq. In this study, we demonstrated TSRdetector’s capability by analyzing normal and diabetic human kidney single-nucleus transcriptomes, to identify kidney cell-type-specific TSR usage and alternative TSR usage associated with diabetes. We also applied TSRdetector to Smart-Seq2 and bulk RNA-seq data to identify the alternative TSR usage during B cell maturation and human ESC transition. Overall, TSRdetector provides an efficient approach for integrating and analyzing single-cell transcriptome data, enabling the detection of cell-type-specific TSR usage in various systems.

## RESULTS

### Workflow of TSRdetector

Non-redundant transcription start regions (TSRs) of a gene were defined as the integrated regions of overlapping first exons from distinct transcripts of the same gene, as described in the Methods section (**Fig. 1a**). The length distribution of TSRs was similar to first exons (**Supplementary Fig. 1a**). In the human genome (CHM13 assembly), 11,121 out of 41,972 RefSeq genes (26%) contain multiple TSRs, while in the mm10 mouse genome, 15,140 out of 54,144 genes (28%) exhibit multiple TSRs. Among these, 82% of multi-TSR genes in humans and 89% in mice encode protein-coding genes (**Fig. 1b**). Unlike single-TSR genes, multi-TSR genes can be transcribed into distinct isoforms depending on the specific TSR utilized, reflecting differences in regulatory mechanisms. To evaluate the 5’-end coverage in single-cell transcriptomic data (scRNA-seq) across different sequencing platforms, we analyzed datasets generated using various technologies. Notably, scRNA-seq data produced using the 10X Genomics 5’ chemistry exhibited significantly higher coverage at the 5’-ends of genes, indicating that this protocol provides superior resolution for distinguishing TSR usage among cell types (**Fig. 1c**). Additionally, Smart-seq and total RNA-seq data demonstrated sufficient coverage of transcript 5’-ends (**Fig. 1c**).

**Fig. 1.**
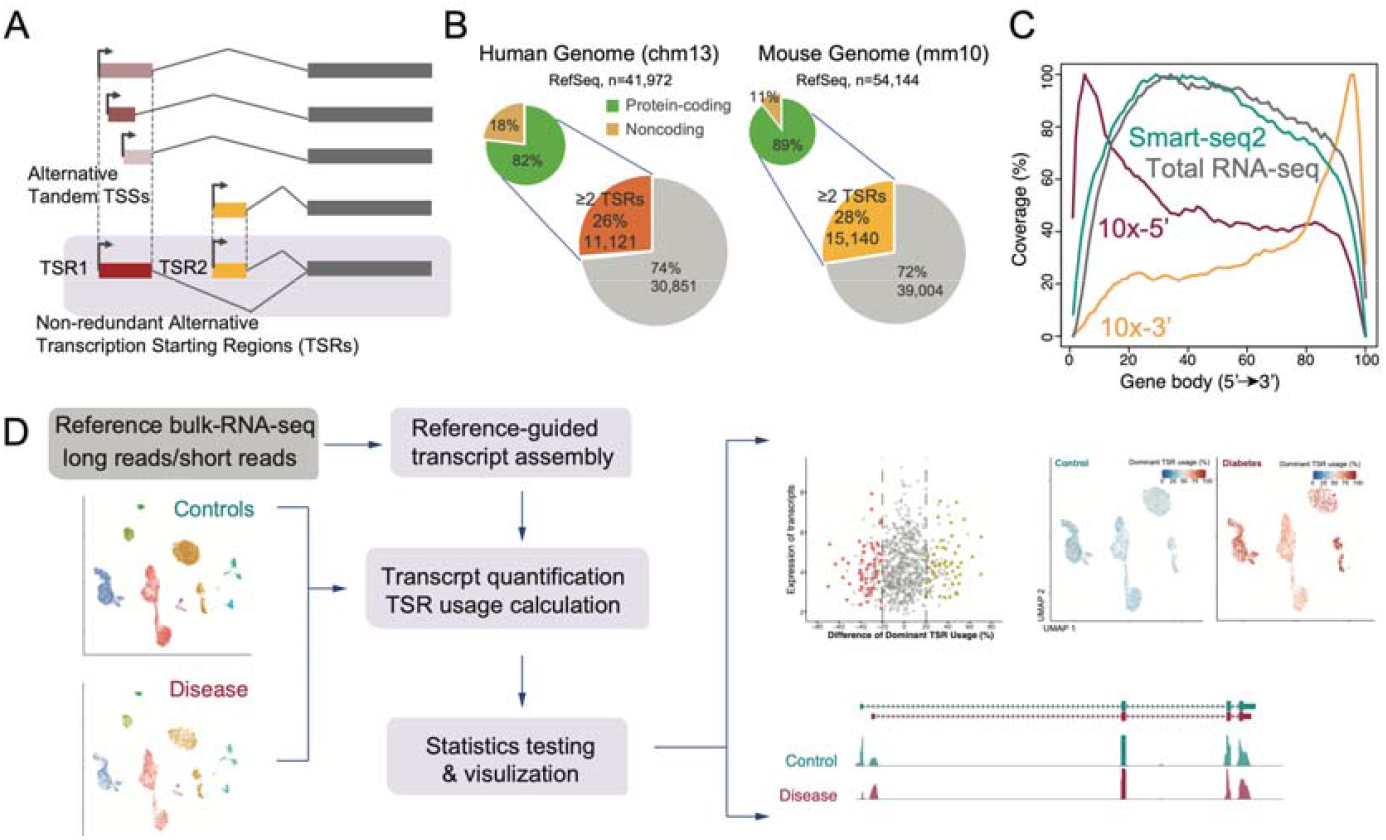
TSRdetector pipeline. **a**, Definition of TSR. TSR is formed by aggregating the genomic coordinates of the first exons from all transcript isoforms belonging to the same gene. **b**, Proportion of unique-TSR and multi-TSRs genes in human and mouse genomes. **c**, The distribution of read coverage across along gene body in 10x Genomics 5’ assay (GSE151302), 10x Genomics 3’ assay (10X PBMC 10K), Smart-seq2 assay (GSE109774), and total RNA-seq assay (GSE175295). **d**, *TSRdetector* workflow: (1) Refernece-guided transcript assembly with long-reads or short-reads RNA-seq; (2) Transcript quantification and TSR usage calculation; (3) Statistics testing and visualization.

To investigate cell-type-specific TSR usage in single-cell transcriptomes, we developed TSRdetector, a computational tool designed to quantify the transcription levels of alternative TSRs using scRNA-seq data (**Fig. 1d; Supplementary Fig. 1b**). To comprehensively capture all potential transcripts, we first designed an annotation-guided long-read sequencing-based transcript assembly module, incorporating long-read RNA-seq data from matched tissues whenever available. Additionally, we processed 54 long-read RNA-seq datasets (**Supplementary table S1**), including 24 PacBio SMRT-seq sequencing, from the ENCODE consortium, along with 30 Oxford Nanopore datasets from a previous study(Gao et al. 2023), to generate comprehensive transcriptome annotations across 52 tissues. Transcripts were assembled by IsoQuant(Prjibelski et al. 2023) with long reads. These three transcriptome annotations were merged into a comprehensive isoform annotation by StringTie(Shumate et al. 2022). These merged annotations are directly available for users to load into TSRdetector (**Supplementary Fig. 1b**).

Using the loaded transcript annotation, TSRdetector aligns scRNA-seq datasets to quantify transcript expression across predefined cell clusters, and calculates TSR usage of all the genes for each cell type (Methods). TSRdetector can perform downstream differential analyses and generate visualization to facilitate the interpretation of TSR dynamics between cell types or different conditions(**Fig. 1d**). In this study, we examined the performance of TSRdetector in different datasets, including single-nucleus RNA-seq (snRNA-seq) generated with 10X Genomics 5’ chemistry(Muto et al. 2021b; Wilson et al. 2019a), Smart-Seq2 data(Schaum et al. 2018; Tabula Muris 2020), and bulk RNA-seq data(Dong et al. 2020).

### Detect cell-type-specific differential TSR usage in human kidney

We applied the TSRdetector to analyze one published snRNA-seq data of five healthy human kidney samples (Muto et al. 2021b; Wilson et al. 2019a), including three male and two female, to investigate differential TSR usage among major kidney cell types. The human kidney is a structurally complex organ composed of multiple cell types, with three major cell types, including distal convoluted tubule principal cells (DCTPC), proximal tubule cells (PT), and loop of Henle cells (LOH), accounting for more than 80% of the total kidney cell population (DCTPC: 31.21%, PT: 30.51%, LOH: 22.30%) (**Fig. 2a**). We noticed that the aggregated sequencing coverage of each cell type, correlated with the proportion of cell-types across all five samples (**Supplementary Fig. S2a**). We focused our analysis on DCTPC and PT cells, which are the most abundant kidney cell types, to investigate how the same gene exhibits distinct TSR usage between these two cell populations.

**Fig. 2.**
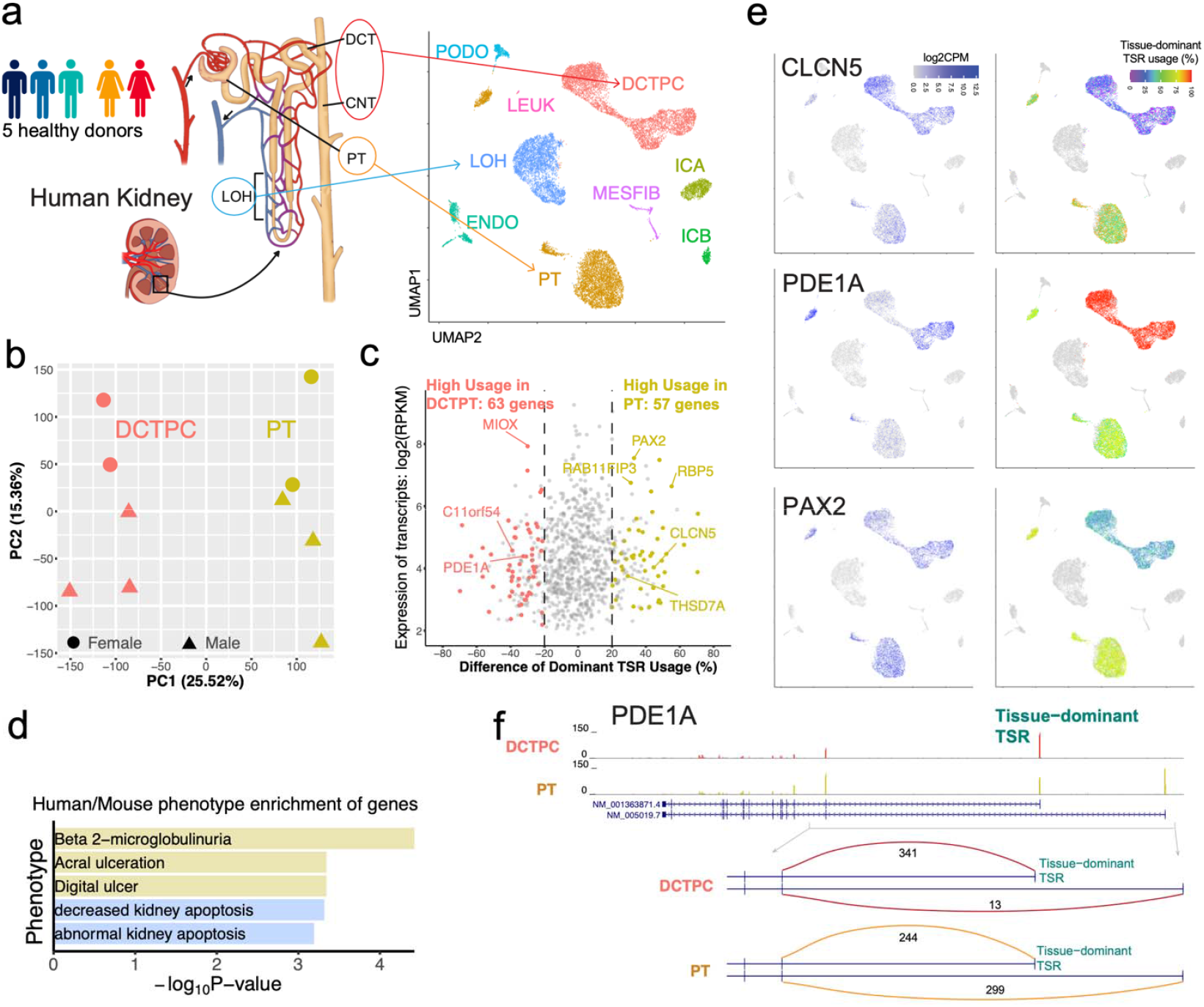
Identification of alternative TSR usage between human kidney DCTPC and PT cells. **a**, snRNA-seq data of Human kidney samples from five healthy donors were obtained, and nine distinct cell type groups were identified and visualized using a UMAP plot (right). **b**, PCA plot of the cell type groups DCTPC and PT from five healthy samples on transcript expression levels. **c**, Comparison of alternative TSR usages of genes between DCTPC and PT. The kidney-specific genes are highlighted. **d**, Gene Ontology enrichment analysis of genes exhibiting significantly different TSR usage between DCTPC and PT cell types. **e**, Gene expression and tissue-dominant TSR usage of *CLCN5, PDE1A*, and *PAX2* in DCTPC and PT cells. **f**, Gene structure, expression, and exon-junction analysis of PDE1A.

TSRdetector first quantified the expression of transcripts in the analyzed samples for selected kidney cell types, and further calculated the TSRs’ usage based on their contribution to overall gene expression (**Supplementary Fig. 1 b**). Principal component analysis (PCA) of transcript expression, revealed a clear separation between DCTPC and PT cells (PC1: 26.04%), while sex differences were captured along PC2 (15.08%) (**Fig. 2b**). In total, TSRdetector identified 120 genes with significant switched usage of TSRs between DCTPC and PT (>20% usage difference, p > 0.05, **Supplementary table S2**). Among these, 63 genes in DCTPC primarily utilized tissue-dominant TSRs, which contribute the most expression of genes at the tissue level. In PT cells, these genes preferentially used alternative secondary TSRs. Conversely, 57 genes in PT cells used tissue-dominant TSRs (**Fig. 2c**). Notably, the TSRs of 96.67% of these genes (116 out of 120) were supported by CAGE-seq data or RAMPAGE data (**Supplementary Fig S2b**).

Further analysis revealed that genes exhibiting alternative TSR usage were enriched in pathways associated with kidney function and disease (**Fig. 2d**). Among the 120 differentially regulated genes, eight were classified as kidney-specific genes according to the Human Protein Atlas database (**Fig. 2c**) [13]. *CLCN5*, which plays a crucial role in trans-4-hydroxy-L-proline catabolism and is considered a potential therapeutic target for primary hyperoxalurias(Solanki et al. 2018), was primarily transcribed (69.8%) from the dominant TSR in PT cells, whereas in DCTPC, this TSR contributed to only 25.2% of transcription (**Fig. 2e**) (**Supplementary Fig.S2c**). *PDE1A*, which encodes a Calcium/Calmodulin-Dependent 3’,5’-Cyclic Nucleotide Phosphodiesterase, has been linked to renal cystic disease and urine-concentrating defects(Wang et al. 2017). *PDE1A* also exhibited cell-type-specific TSR usage: in DCTPC, 99.2% of transcription was initiated from the dominant TSR, while in PT cells, only 68.4% of transcription originated from this TSR (**Fig. 2e,f**; **Supplementary Fig.S2c**). *PAX2*, a key regulator of urine concentration(Laszczyk et al. 2020), exhibited distinct TSR usage patterns, with its tissue-dominant TSR predominantly utilized in PT cells but not in DCTPC (**Fig. 2e)**.

We further examined the general expression levels of the 120 genes that exhibited differential dominant TSR usage between DCTPC and PT cells. Surprisingly, 79 of these genes were expressed at similar levels in both cell types, indicating that changes in TSR usage did not necessarily correlate with overall gene expression. Only 22 genes and 19 genes showed significant changes in gene expression levels alongside differential TSR usage in DCTPC and PT cells, respectively (**Fig. 3a; Supplementary table S3**). For example, *ARID5B*, a transcriptional coactivator whose homozygous loss causes kidney defects in mice, binds to the 5’-AATA[CT]-3’ core sequence and recruits phosphorylated *PHF2* to mediate H3K9me2 demethylation during tissue development and carcinogenesis(Baba et al. 2011). Although *ARID5B* gene expression levels were similar between DCTPC and PT cells, the tissue-dominant TSR was preferentially used in PT cells (81.1%) but only in 22.0% of cases in DCTPC (**Fig. 3b, 3c**). Exon junction alignment of RNA-seq reads further confirmed that PT cells predominantly utilized the dominant TSR, driving over 80% of *ARID5B* expression, whereas DCTPC cells relied on a tissue-secondary TSR, contributing to approximately 75% of *ARID5B* expression (**Fig. 3d**).

**Fig. 3.**
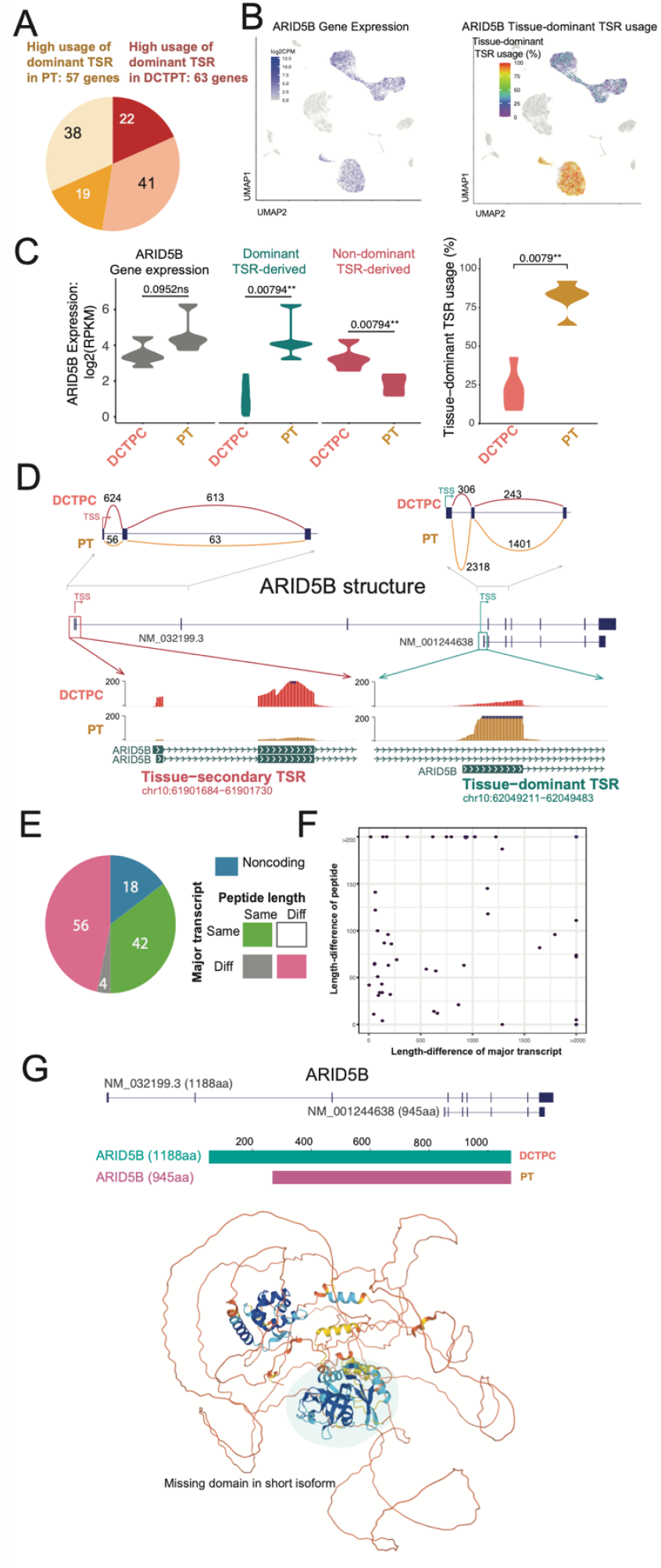
Alternative TSR usage induces protein isoform switching without altering overall gene expression. **a**, 22 (dark red) and 19 (orange) genes with significantly alternative TSRs usage between DCTPC and PC cells are differentially expressed genes. **b**, Gene expression (left) and tissue-dominant TSR usage (right) of *ARID5B* in the UMAP plot. **c**. Left: expression of *ARID5B* gene, its dominant TSR-derived transcript, and non-dominant TSR-derived transcript in DCTPC and PT cells. Right: Tissue-dominant TSR usage of *ARID5B*. **d**. Exon-junction analysis and gene expression of *ARID5B*. **e**. Alternative TSR usage in 56 of 120 genes leads to the production of protein isoforms with different lengths between DCTPC and PT cells. **f**. The length differences of the major transcript isoforms between DCTPC and PT cells do not correlate with the length differences of their resulting peptide products. **g**. Gene structure of *ARID5B* and predicted 3D structure of the long peptide isoform (AlphaFold2).

Among the 120 genes with differential tissue-dominant SR usage between DCTPC and PT cells, 78 genes exhibited changes in their major transcribed isoform (defined as an isoform contributing to >50% of the gene’s total expression) (**Supplementary table S4**). Notably, 72% of these genes underwent isoform switching, resulting in predicted alterations in translated peptides (**Fig. 3e**). Interestingly, the changes in peptide length were not correlated with changes in RNA transcript length (**Fig. 3f**). In DCTPC cells, *ARID5B* transcription was primarily driven by the tissue-dominant TSR, leading to the production of the full-length protein product (1188 amino acids). However, in PT cells, transcription was mainly initiated from a tissue-secondary TSR, producing a C-terminal truncated protein (945 amino acids). AlphaFold structural predictions suggested that this truncated isoform lacked the well-organized C-terminal coiled-coil and beta-hairpin motif, which are key structural elements of the full-length protein (**Fig. 3g**).

### Transposable elements derived transcription in major human kidney cell types

Transposable elements (TEs) are highly repetitive DNA sequences that are widely distributed across mammalian genomes. It is well established that TEs can initiate gene transcription and contribute to tissue-specific gene regulation. To investigate the role of TE-derived transcription in human kidney cell types, we utilized TSRdetector to analyze TE-derived transcription start regions (TE-TSRs) across different kidney cell types. In total across the three major kidney cell types, we identified 1,169 genes that initiate transcription from TE-derived TSRs, which can contribute over 10% of the transcription of overall gene expression (**Supplementary table S5**). Among these, 714 genes were commonly detected in all three cell types, while 56, 118, and 48 genes exhibited cell-type-specific TE-derived expression in DCTPC, LOH, and PT cells, respectively (**Fig. 4A**). This suggests that TE-derived transcription is a prevalent regulatory mechanism across in the human kidney.

**Fig. 4.**
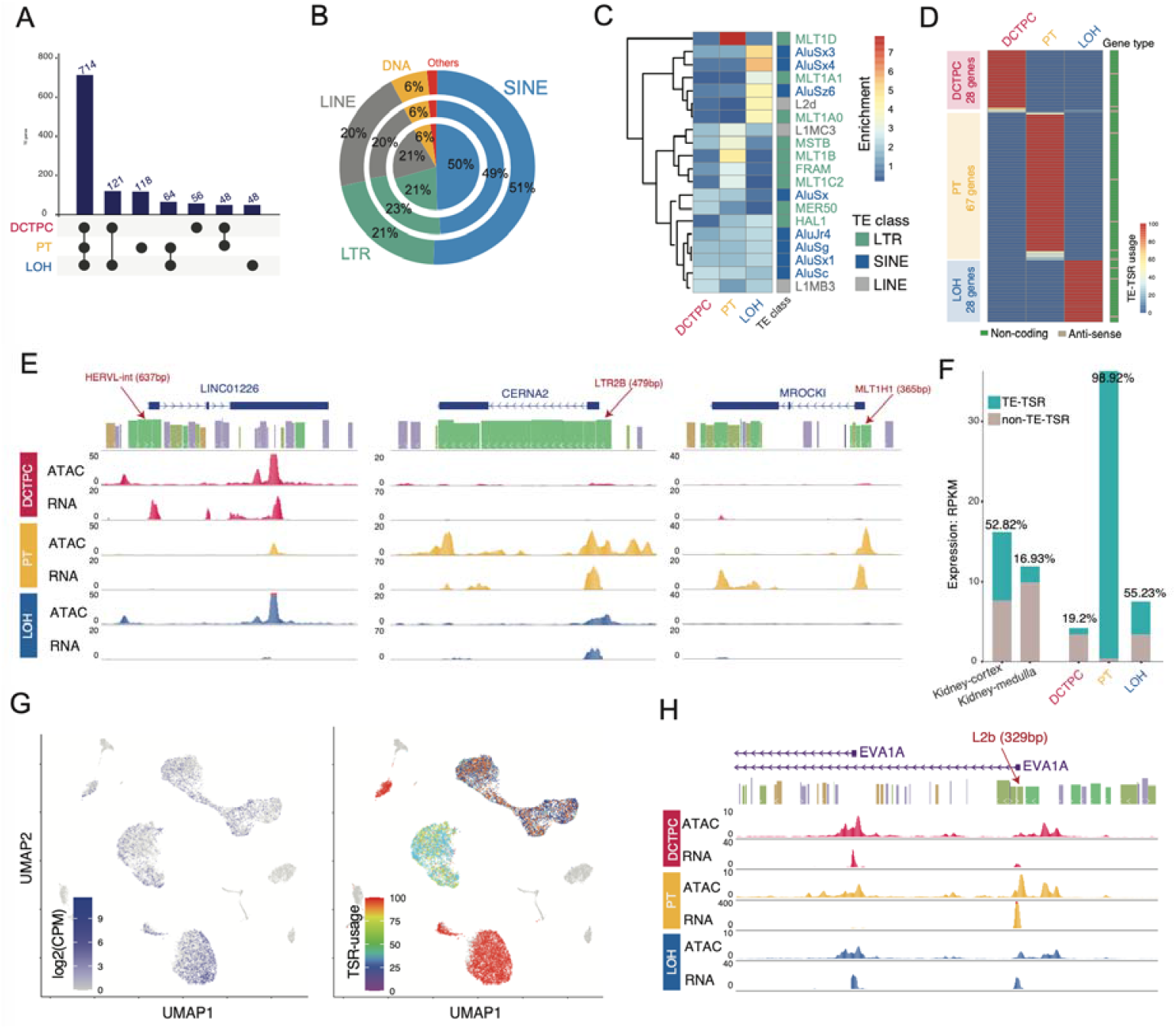
Transposable element-derived TSRs in human kidney cell types. **a**. UpSet plot of 1,169 genes that initiate transcription from TE-derived TSRs across the three major kidney cell types. **b**. TE classes contributing to TE-derived TSRs across the three major kidney cell types, shown from outer to inner rings: DCTPC, PT, and LOH. **c**. TE subfamily enrichment of TE-TSRs in three major kidney cell types. **d**. 123 TE-initiated cell-type-specific non-coding or anti-sense genes. **e**. Examples of TE-initiated cell-type-specific genes. **f**. Proportional usage of the L2b-derived TSR of *EVA1A* in human kidney tissues and across the three major kidney cell types. **g**. Gene expression level and L2b-derived TSR usage of *EVA1A*. **h**. L2b-derived TSR of *EVA1A* is highly accessible (ATAC-seq) in PT cells.

Further analysis revealed that SINE elements contributed over 50% of TE-TSRs in kidney cell types (**Fig. 4B**), a proportion significantly higher than the genome-wide background (**Supplementary Fig. S3a**). TE subfamily enrichment analysis demonstrated distinct cell-type-specific activation of specific TE subfamilies. For example, AluSx3, AluSx4, and AluSz6 were highly enriched in LOH cells, whereas MLT1D and MLT1B were predominantly activated in PT cells (**Fig. 4C**). Among the 222 cell-type-specific TE-initiated genes, 123 genes were classified as non-coding genes (**Fig. 4D**), indicating that cell-type-specific TE activation contributes significantly to non-coding RNA transcription. Additionally, 173 TE-initiated genes exhibited significant differential TE-TSR usage (>40% usage difference) between at least two cell types (DCTPC/PT/LOH: 37/97/39) (**Supplementary Fig. S3b; Supplementary table S6**).

One notable example is *EVA1A*, a key regulator of autophagy and apoptosis, which is highly expressed in the liver and kidney. Bulk RNA-seq data from GTEx indicated that an L2b TE-derived TSR contributes to ∼52% of EVA1A transcription in the kidney cortex and ∼17% in the medulla (**Fig. 4E**). TSRdetector analysis further revealed that this L2b-TSR exhibits striking cell-type specificity: in PT cells, it initiates ∼99% of *EVA1A* expression, whereas in DCTPC cells, it drives only ∼19% of total EVA1A transcription (**Fig. 4F, G; Supplementary Fig. S3c, d**). Furthermore, single-nucleus ATAC-seq data demonstrated significantly stronger open-chromatin signals at the L2b-TSR in PT cells, compared to DCTPC and LOH cells (**Fig. 4H**). This suggests that chromatin accessibility at TE-TSRs plays a critical role in cell-type-specific transcriptional regulation in the kidney.

### Switched TSRs usage in diabetic kidneys

TSR usage can be influenced by various cell states, including differentiation, disease progression, and responses to external stimuli. To investigate diabetes-induced changes in TSR usage within the three major kidney cell types, we applied TSRdetector to compare diabetic kidneys with healthy controls. Cell-type alignment analysis revealed that diabetic kidney cells mapped well to their healthy counterparts, suggesting minor global expression changes in diabetic kidneys (**Fig. 5a**). However, TSRdetector identified significantnly alterations of TSR usage: 55 genes in DCTPC, 49 genes in PT, and 65 genes in LOH (**Fig. 5b; Supplementary Fig. S4a; Supplementary table S7**). Among these, six kidney-specific genes, *KCNJ1, TMEM37, ASPA, POU5F1, PAX2*, and *MYL12B*, exhibited significant alternative TSR usage in diabetic kidneys. Gene Ontology (GO) enrichment analysis of these genes indicated enrichment in several key biological processes relevant to kidney function and disease, such as intracellular transport, DNA damage, and cell cycle checkpoint (**Fig. 5c**).

**Fig. 5.**
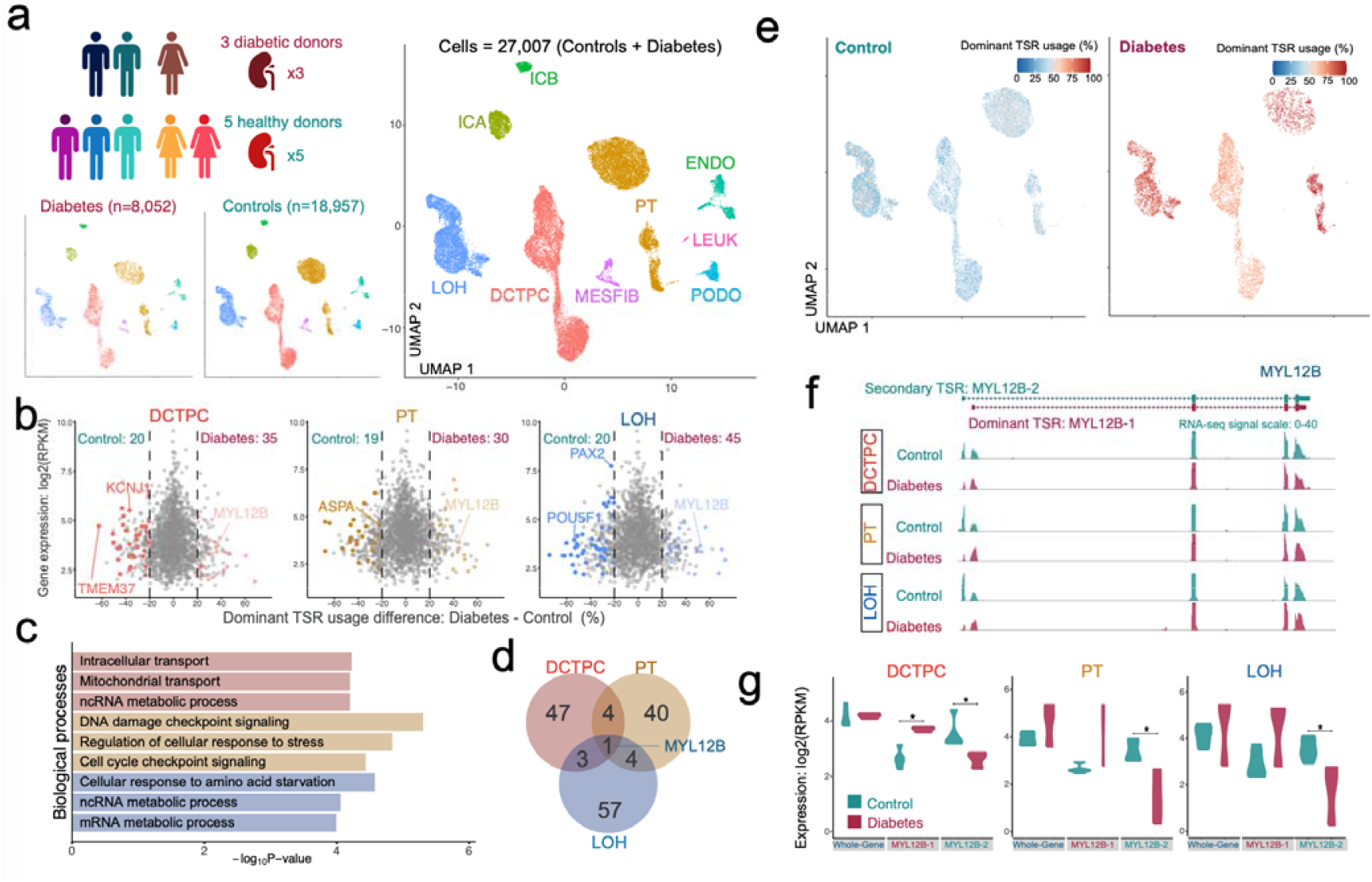
Diabetes-induced alternative TSR usage in human kidney. **a**, snRNA-seq data of human kidney samples from five healthy and three diabetic donors were sequenced. UMAP plots show the nine identified cell type groups in diabetic samples (left), control samples (middle), and combined (right). **b**. Alternative usage of tissue-dominant TSRs between control and diabetic conditions in three cell type groups. **c**. Enriched biological processes GO terms of genes with significantly altered tissue-dominant TSR usages between control and diabetic conditions. **d**. Overlapped genes with significantly altered tissue-dominant TSR usages in three cell type groups. **e**. Dominant TSR usage of *MYL12B* in control and diabetic kidney. **f**. Aggregated snRNA-seq single of *MYL12B* of three cell types in control and diabetic kidneys. **g**. Expression of *MYL12B* gene, its dominant TSR-derived transcript, and non-dominant TSR-derived transcript in three kidney cell groups.

Interestingly, most genes exhibited altered TSR usage in a cell-type-specific manner, except for *MYL12B*, which showed differential TSR usage across all three kidney cell types (**Fig. 5d**). *MYL12B*, a myosin regulatory subunit, plays a crucial role in controlling cell contractility via phosphorylation, and elevated phosphorylated *MYL12B* levels in plasma have been identified as an early biomarker for acute kidney injury(Wu et al. 2015). In all three kidney cell types, we observed significant differences in TSR usage between healthy and diabetic kidneys (**Fig. 5e, 5f**). However, the overall expression levels of *MYL12B* remained stable (**Fig. 5g**), suggesting that diabetic conditions alter the transcriptional regulation of *MYL12B* isoforms without affecting total gene expression levels.

### Identification of altered TSR usage during B cell maturation by using SMART-seq2

To demonstrate the capability of TSRdetector in detecting TSR usage across different sequencing technologies, we applied it to analyze a SMART-seq2 time-course dataset tracking B cell maturation (**Fig. 6A**). Re-analysis of the SMART-seq2 dataset, which comprises 1,553 single cells, revealed that the three stages of B cell development were distinctly separated along a continuous trajectory, reflecting the progression of B-cell maturation (**Fig. 6B**). Using TSRdetector, we identified 54 genes that exhibited significant shifts in TSR usage across different maturation stages (**Fig. 6C; Supplemental table S8; Supplemental Fig S5a, b**). Many of these genes were functionally linked to DNA damage response, lymphocyte polarity regulation, and immune signaling pathways, suggesting that alternative TSR usage plays a role in transcriptional adaptation during B cell development (**Fig. 6D; Supplemental Fig S5c**).

**Fig. 6.**
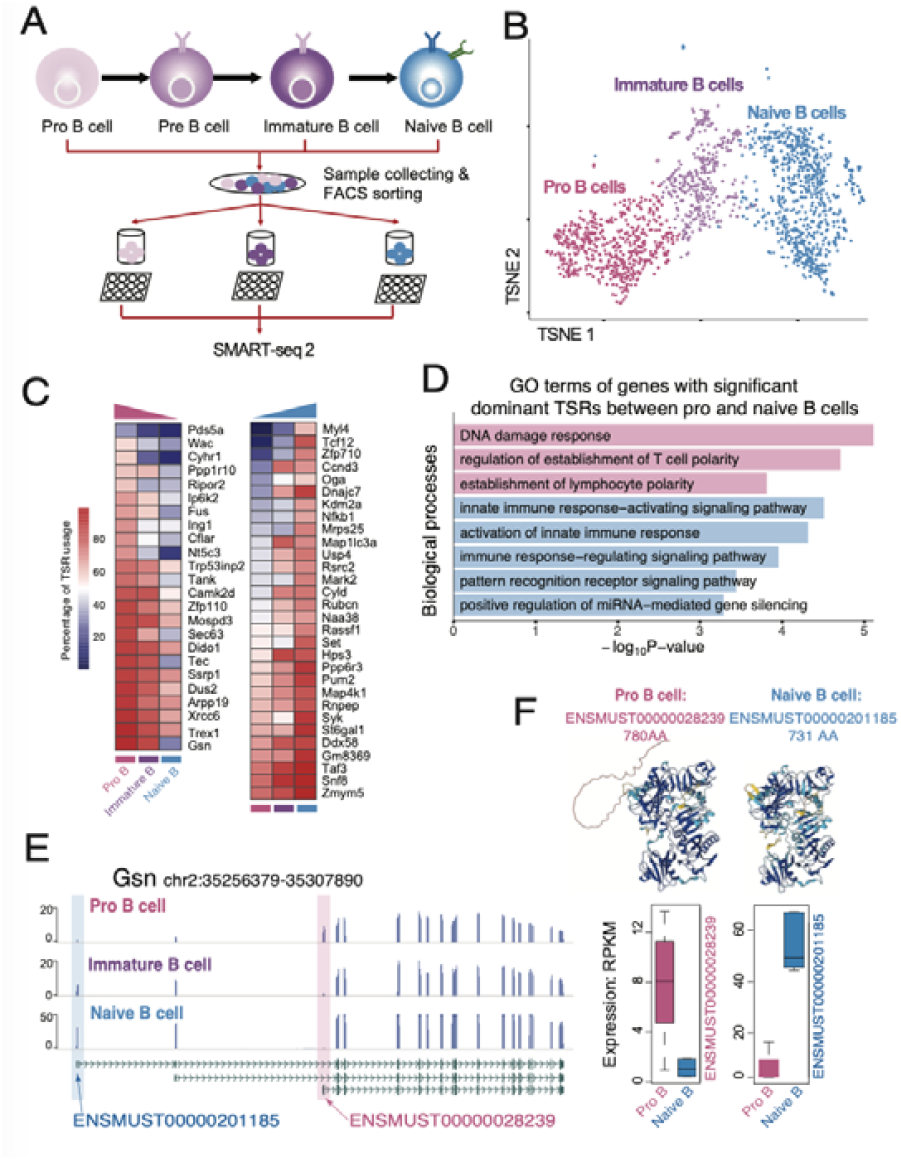
TSRdetector identified alternative TSR usage during mouse B-cell maturation. **a**, Mouse B-cell development and experiment design. **b**, t-SNE plots of Pro B cells, immature B cells, and Naïve B cells. **c**, Heatmap plots of significantly altered tissue-dominant TSR usages during B-cell maturation. **d**, Enriched biological processes (GO terms, q-value <0.1) of genes with significantly altered tissue-dominant TSR usages between pro B cells and naïve B cells. **e**, Aggregated SMART-seq2 single of *Gsn* during B-cell maturation. **f**. Predicted 3D structure of the long and short Gsn peptide isoform (AlphaFold2) and switched expression during B-cell maturation.

One particularly notable case is *Gsn* (Gelsolin), an actin-modulating protein implicated in cytoskeletal remodeling, immune response, and amyloidosis pathology(Piktel et al. 2018). TSRdetector revealed that Gsn utilizes two distinct TSRs to transcribe different isoforms in Pro B cells and naïve B cells (**Fig. 6E**). In Pro B cells, the longer isoform (ENSMUST00000028239, 780 amino acids) was predominantly expressed. However, during the transition to naïve B cells, TSR usage shifted, resulting in the preferential expression of a shorter isoform (ENSMUST00000201185, 731 amino acids) (**Fig. 6F**). Meanwhile, Tcf12, a main helix-loop-helix transcription factors that regulates B cell and T cell development, also indicated strong alternative TSR switch. Tcf12 continuously downregulates the expression of the short isoform during B cell maturation but upregulates the expression of the long isoform (**Supplemental Fig S5e**). Such results suggest that alternative TSR selection during B cell maturation contributes to isoform switching, and can further affect protein function and cellular behavior.

### Identification of altered TSR usage in hESC cell states transition

Bulk RNA-seq is the most widely used method for cost-efficient transcriptome profiling, offering excellent gene-body coverage and diverse transcripts profiling (**Fig. 1C**). To evaluate the performance of TSRdetector in detecting alternative TSR usage in bulk RNA-seq data, we applied it by comparing human primed and naïve stem cells. Human naïve stem cells can be derived from primed stem cells through treatment with the 5i/L/A cocktail, which induces a global transcriptomic shift and reprograms the cells into a distinct pluripotent state(Huang et al. 2021) (**Fig. 7A**). Principal component analysis (PCA) confirmed that this transition significantly alters gene expression patterns, distinguishing naïve and primed states at the transcriptome-wide level (**Fig. 7B**). Using TSRdetector, we identified 137 genes that exhibited significant changes in TSR usage during this transition (**Fig. 7C, Supplementary table S9**), including key regulators of early embryonic development.

**Fig. 7.**
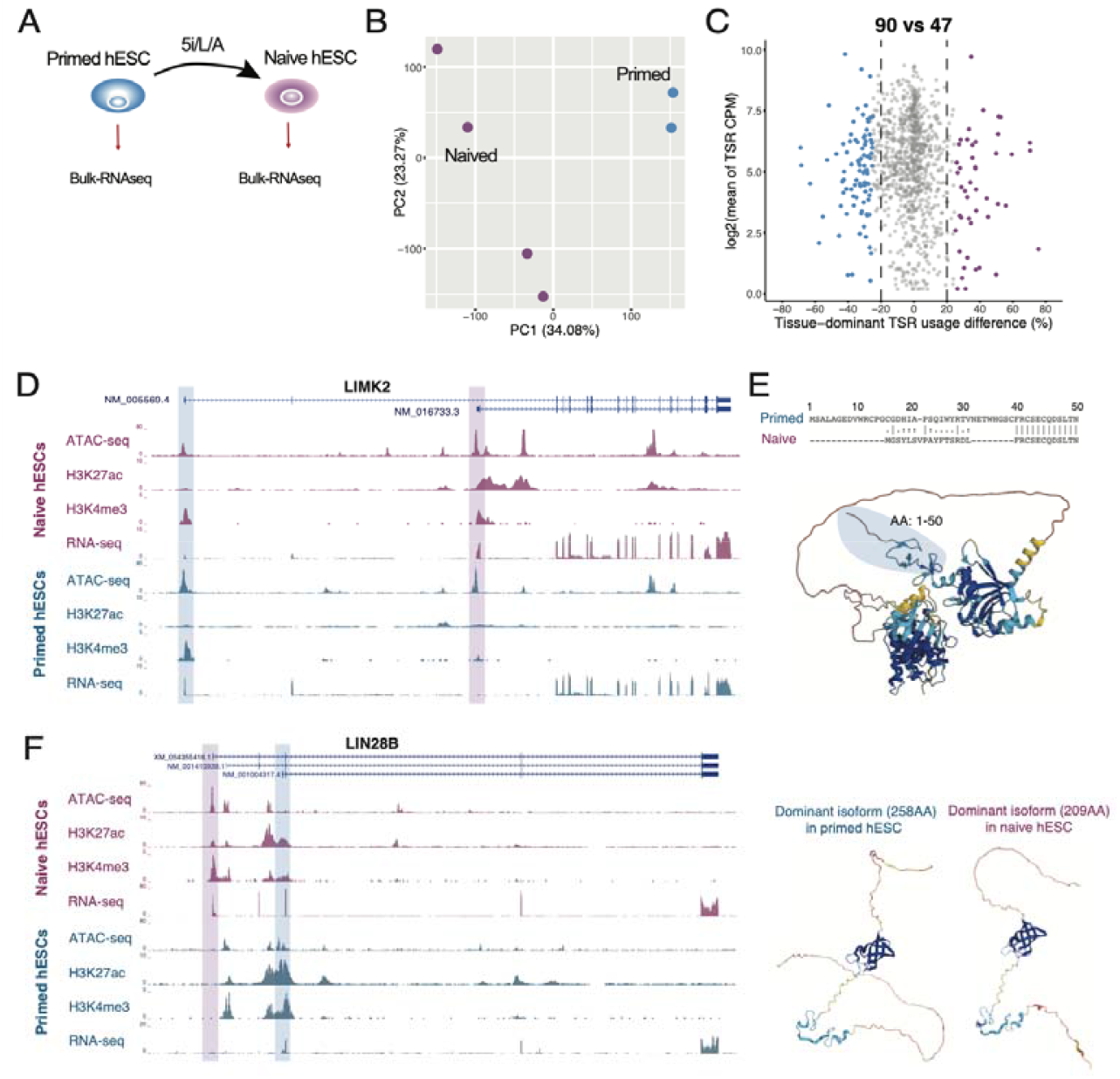
TSRdetector identified alternative TSR usage during hESC cell states transition. **a**. Naïve hESC is derived from primed hESC with the 5i/L/A cocktail. **b**. PCA plot of naïve and primed hESC samples. **c**. Genes with significant TSR alteration between primed and naïve hESCs. **d**. Expression (RNA-seq) and epigenetic modification (ATAC-seq and H3K27ac and H3K4me3 ChIP-seq) of LIMK2 in naïve and primed hESC. **e**. Predicted 3D structure of alternative peptide isoforms of LIMK2 (AlphaFold2). **f**. Expression and epigenetic modifications of LIN28B(left) and predicted 3D structure of alternative peptide isoforms (right).

One notable example is *LIMK2*, a serine/threonine protein kinase that plays a critical role in actin filament regulation(Maekawa et al. 1999) and is directly involved in cell division and motility(Tersoff 1992). In primed hESCs, only the long isoform of *LIMK2* was expressed, whereas naïve hESCs predominantly utilized an alternative TSR, resulting in the dominant expression of a shorter isoform. This TSR switch was accompanied by epigenetic remodeling in the regulatory landscape surrounding the two TSRs (**Fig. 7D**). The isoform switch also led to a truncated *LIMK2* protein in naïve hESCs, potentially affecting its function in cytoskeletal dynamics and cell migration (**Fig. 7E**). Similarly, *LIN28B*, an RNA-binding protein essential for the naïve-to-primed pluripotency transition(Zhang et al. 2016), demonstrated TSR-dependent isoform switching. In primed hESCs, *LIN28B* transcription was initiated primarily from an intragenic AluJB-derived TSR, producing a longer isoform (258 amino acids). Upon reprogramming to the naïve state, an alternative upstream 5’ TSR was activated, leading to the expression of a shorter *LIN28B* isoform (209 amino acids) (**Fig. 7F**). This shift suggests a potential functional adaptation of *LIN28B* in the naïve state, possibly influencing RNA metabolism and pluripotency regulation.

## DISCUSSION

Gene transcription regulation starts from activation of specific promoters at precise times in a highly cell-type-specific manner. Therefore, accurately measuring the usage of transcription start regions (TSRs) for each transcript and assessing their regulatory impact on overall gene expression is crucial. Here, we present TSRdetector, an integrative package designed to accurately quantify TSR usage and isoform expression at the single-cell level. TSRdetector is optimized for processing single-cell RNA-seq data that contains 5’ end or full-length transcript information while also supporting bulk RNA-seq data. To demonstrate its power and utility, we applied TSRdetector to multiple datasets, including 10X Genomics snRNA-seq (5’-end protocol), SMART-seq2, and bulk RNA-seq. Across all tests, TSRdetector effectively identified differential TSR usage among various cell types, during cell state transitions, and in human disease contexts. These results highlight its potential for advancing our understanding of transcription regulation in diverse biological settings.

In the human kidney snRNA-seq dataset (10X, 5’ protocol), TSRdetector revealed cell-type-specific TSR usage in hundreds of genes between DCTPC and PT cells, including kidney-specific genes like *PAX2, CLCN5*, and *PDE1A*. Despite similar gene expression levels, many genes showed distinct TSR usage, highlighting a limitation of traditional differential expression analysis. Nearly half of these TSR shifts resulted in protein isoforms of different lengths—independent of mRNA length—suggesting that alternative TSR usage is a key regulatory mechanism. For example, *ARID5B* used a longer isoform in DCTPC and a shorter, truncated isoform in PT cells, potentially affecting protein function and localization.

We also utilized TSRdetector to investigate transposable element (TE)-derived TSRs in human kidney cells. Although nearly half of the human genome originates from TEs, the majority remain highly suppressed through mechanisms such as epigenetic modifications and ZNF-mediated silencing, except for a small subset that has been domesticated as regulatory elements, including promoters and enhancers. In our study, 1,169 genes were found to initiate transcription using TE-derived TSRs across all three kidney cell types, with 714 of these TSRs being shared among them. Notably, ∼90% of the 222 cell-type-specific TE-TSRs were associated with long noncoding RNAs (lncRNAs), supporting previous findings that TEs contribute significantly to species-specific lincRNAs(Kapusta et al. 2013). Additionally, we observed that SINE-derived TSRs were highly active in transcription initiation, particularly the AluSx3 and AluSx4 elements, which were strongly activated in LOH cells. This suggests a potential unique activation mechanism for Alu elements in human kidney function and regulation.

In human diabetic kidney, TSRdetector identified approximately 150 genes with significant changes in TSR usage compared to healthy controls. These genes were enriched in biological processes strongly linked to diabetes progression, including mitochondrial function, DNA damage response, cell cycle checkpoint regulation, and cellular stress response. Notably, *MYL12B*, an early biomarker for acute kidney injury(Wu et al. 2015), exhibited altered TSR usage across all three kidney cell types, suggesting a potential link between kidney injury and diabetes progression. While previous studies have shown that diabetes can cause subtle disruptions in gene regulation, our findings reveal an alternative regulatory mechanism in diabetic kidney disease, highlighting TSR dynamics as a potential contributor to gene expression changes in this chronic condition.

We also observed the strong performance of TSRdetector in SMART-seq2 and bulk RNA-seq data. Both data types contain sufficient 5’ end transcript information, enabling TSRdetector to efficiently identify differential TSR usage during cell fate transitions of B cell maturation and human embryonic stem cells (hESCs). Notably, we identified LIN28B, a key regulator in naïve hESCs, as being specifically transcribed through an AluJB-derived TSR, highlighting the potential role of transposable element (TE) activation in human embryonic development. This finding underscores the significance of TE-derived regulatory elements in shaping cell identity and differentiation.

Across multiple tests, TSRdetector demonstrated a strong ability to evaluate TSR usage across different data types, including single-cell RNA-seq and bulk RNA-seq. Its flexibility allows it to be widely applied in various studies, providing deeper insights into cell-type-specific TSR usage and improving our understanding of gene regulation. However, TSRdetector has certain limitations. Notably, it does not support the widely used 3’ end single-cell RNA-seq protocols, as these datasets lack sufficient information to characterize 5’ transcript ends. Additionally, TSRdetector currently provides TSR usage estimates at the cell-type level, rather than true single-cell resolution, due to sequencing depth constraints. Despite these limitations, TSRdetector remains a comprehensive and adaptable tool for analyzing TSR usage at the single-cell-type level and is compatible with most RNA-seq datasets containing 5’ end information.

## METHODS

### TSRdetector pipeline

#### Reference-guided Transcript assembly

TSRdetector is compatible with pre-existing transcript annotations, including RefSeq, GENCODE, and Ensembl. Additionally, it offers a comprehensive transcript annotation derived from long-read RNA-seq data collected from 52 human tissues (**Supplementary table S1**). For users interested in generating custom transcript annotations, TSRdetector supports reference-guided transcriptome assembly using either short-read or long-read RNA-seq data from the analyzed samples. It includes built-in modules for transcript assembly: StringTie(Shumate et al. 2022) is used for assembling short-read RNA-seq data, while IsoQuant (Prjibelski et al. 2023)is employed for assembling long-read RNA-seq data. Long-read RNA-seq data are first aligned to the reference genome using minimap2(H. Li 2021). These raw alignments are then refined using TranscriptClean(Wyman and Mortazavi 2019), which corrects splice junctions by incorporating short-read alignment data when available. Additionally, unaligned long reads are error-corrected using LoRDEC(Salmela and Rivals 2014), which utilizes the reference genome and, when applicable, aligned short-read RNA-seq data to enhance alignment accuracy and sensitivity.

*Definition of reference TSRs*

Using the loaded reference transcript annotation, TSRdetector defines transcription start regions (TSRs) for each gene by aggregating the genomic coordinates of the first exons from all transcript isoforms belonging to the same gene. Redundant regions are merged to create a set of non-overlapping TSRs per gene. When a gene has multiple TSRs, they are sequentially ordered based on their genomic positions, taking the transcriptional strand into account to preserve the correct directionality (Fig. 1b). TSRs that overlap but originate from different genes are treated as distinct entities, with boundaries assigned according to their respective gene annotations.

#### Identification and quantification of TSRs in assayed samples

After trimming adapters, TSRdetector first aggregates the reads of scRNA-seq data for each cell types based on input cell type barcode/clustering information. TSRdetector aligns all aggerated reads of each cell type in each sample to the reference genome by using STAR (Dobin et al. 2013), and the transcript expression levels are estimated by using RSEM (B. Li and Dewey 2011) based on the loaded transcripts annotation. The usage of each TSR (*U*_*TSR_i*_)is quantified by calculating the proportion of the total gene expression contributed by transcripts originating from that specific TSR, as follows:

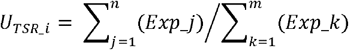

In which *Exp_j* represents the esssxpression of transcript *j* that derived from *TSR_i* and *n* is total number of transcripts initiated from *TSR_i. Exp_k* represents the expression of transcript *k* from all transcripts (*m*)of same gene. TSRdetector will organize the TSR usage table after collecting the usage of TSRs for user selected cell types in all samples, and only keeps high quality TSRs based on expression (RPKM of TSR > 1) and mapping criteria (uniquely aligned splicing reads >5) to downstream analysis.

#### Detection of significantly changed dominant TSRs between conditions

For downstream analysis, TSRdetector selects genes that have at least two distinct transcription start regions (TSRs). The usage percentages of each TSR are grouped according to experimental or biological conditions (e.g., treatment vs. control). A Wilcoxon rank-sum test is then applied to compare TSR usage distributions between conditions. Genes with a resulting p-value below a defined threshold (e.g., 0.05) and a difference in the mean usage of the tissue-dominant TSR greater than 20% between groups are considered to exhibit significant differential usage of TSRs.

*Data visualization*

TSRdetector generates a BED-format file containing genomic coordinates of significantly differentially used TSRs. This file can be directly loaded into the Epigenome Browser (D. F. Li et al. 2022) for visualization of the affected regions. In addition to the BED file, the pipeline produces stacked percentage bar plots and sashimi plots for each gene exhibiting significant TSR usage differences. The bar plots, illustrating TSR usage distribution across conditions, are generated using the ggplot2 R package(Ginestet 2011). Sashimi plots are produced with ggsashimi(Garrido-Martin et al. 2018), using the settings *-S strand -M 3 --base-size=20 -C 3 -O 3--shrink --alpha 0*.*5*, which require a minimum of three supporting reads per visible splice junction. Only strand-specific reads aligned to the same strand as the gene locus are included in the visualization

### Human kidney single-nucleus RNA-seq and single-nucleus ATAC-seq data processing

The single-nucleus RNA sequencing (snRNA-seq) datasets used in this study were derived from kidney samples of eight individuals, including five healthy controls (three males and two females) and three patients diagnosed with diabetic kidney disease (two males and one female), as previously described in previous studies (Muto et al. 2021a; Wilson et al. 2019b). To reduce technical noise and facilitate interpretation, cell types with similar transcriptional signatures and anatomical proximity were clustered into nine broader cell-type groups. These groupings were based on established renal cell type classifications and spatial organization within the nephron and surrounding compartments. Specifically: DCTPC includes Distal Convoluted Tubule 1 and 2 (DCT1, DCT2) and Principal Cells (PC), which are functionally associated with distal tubular processing; PT includes Proximal Convoluted Tubule (PCT), Proximal Epithelial Tubule (PET), Proximal Straight Tubule (PST), and Proximal Tubule VCAM1+ (PTVCAM1), representing the reabsorptive segments of the proximal nephron; LOH includes Thick Ascending Limb (TAL), Cortical TAL (CTAL), and Medullary TAL (MTAL), key segments in the loop of Henle involved in salt reabsorption; ICA represents Intercalated Cells A, which regulate acid-base balance; ENDO includes Glomerular Endothelial Cells (GEC) and Peritubular Capillary Cells (PTC), both critical components of the renal vascular system; PODO includes Podocytes, specialized epithelial cells that form the filtration barrier in glomeruli; MESFIB includes Mesangial Cells (MES) and Fibroblasts (FIB), which contribute to structural support and extracellular matrix production; ICB includes Intercalated Cells B, involved in bicarbonate secretion and pH regulation; LEUK comprises Leukocytes, including immune cells infiltrating the kidney tissue.

TSRdetector processed all human kidney single-nucleus RNA-seq datasets using the T2T-CHM13 v2.0 reference genome assembly. Transcript expression levels were quantified, and TSR usage was calculated based on RefSeq transcript annotations, which were obtained from the UCSC Genome Browser. Specially, the three most prevalent cell-type groups across all samples were DCTPC, PT, and LOH, reflecting their abundant representation in renal cortical tissue (Fig. S2a). Differential TSR analysis were performed between DCTPC and PT cell groups in healthy kidneys, and between healthy kidneys and diabetic kidneys in DCTPC, PT, and LOH cell groups. The transposable elements (TE) annotation file of T2T CHM13v2.0 genome assembly were obtained from the hg38 annotation through liftOver, and the TE-derived TSRs were defined as the 5’ end of TSRs located within TEs.

For differential gene expression (DEG) analysis, clustered single-nucleus RNA-seq reads were aligned to the T2T-CHM13 v2.0 reference genome using STAR aligner (version 2.5.4b). Gene-level quantification was performed with featureCounts from the Subread package (version 1.4.6)(Liao et al. 2014), utilizing the corresponding T2T-CHM13 v2.0 gene annotation. To correct for unwanted sources of variation in gene expression data, the RUVr function from the RUVSeq (Risso et al. 2014) normalization package was applied. The number of factors of unwanted variation (k) was empirically set to 2 for comparisons between DCTPC and PT groups in healthy control samples, and to 3 for comparisons between control and diabetic samples within each of the three major cell type groups (DCTPC, PT, and LOH).

Normalized gene counts were subsequently imported into DESeq2(Love et al. 2014) for DEG analysis. Only genes with counts per million (CPM) greater than 1 in at least one condition were retained for downstream analysis. The filtered count matrix was transformed using the regularized log transformation (rlog) provided by DESeq2 to stabilize variance across samples. Statistical significance of differential expression was assessed using the Wald test implemented in DESeq2, with multiple testing correction performed via the Benjamini-Hochberg method using p.adjust function in R. Genes were considered differentially expressed if they met the criteria of an absolute log2 fold change > 1 and a false discovery rate (adjusted p-value) < 0.01.

Single-nucleus ATAC-seq data were downloaded from GSE195460, and reads were clustered by barcode to identify cell types, consistent with previous classifications. The clustered raw reads were aligned to the T2T-CHM13 v2.0 reference genome and processed using the AIAP pipeline(Liu et al. 2021), which includes four main steps: data processing, quality control, integrative analysis, and data visualization. Narrow peak files from all ATAC-seq libraries were merged using the merge function from the BEDTools suite, and peak-level read counts were quantified using the coverage command in BEDTools.

### SMART-seq2 data processing

The mouse bone marrow B cells SMART-seq2 datasets were obtained from the Tabula Muris Consortium, originally generated using fluorescence-activated cell sorting (FACS)(Schaum et al. 2018; Tabula Muris 2020). The SMART-seq2 B cells datasets contain three distinct developmental stages: pro-B (n = 517), immature B (n = 344), and naïve B (n = 692). To ensure sufficient transcript coverage, cells from each developmental stage were randomly grouped into clusters with similar total read counts, resulting in six biological replicates per stage (Fig. S5a), and to facilitate the downstream differential analysis. TSRdetector was applied using Ensembl 98 gene annotations (GRCm38 reference genome) to quantify transcript-level expression and calculate the transcription start region (TSR) usage. To identify significantly differentially used TSRs between the naïve and pro-B stages, a Wilcoxon rank-sum test was performed with a p-value threshold of 0.05 and a minimum usage ratio difference of 20%. In parallel, differential gene expression (DEG) analysis between pro-B and naïve B cells was conducted using DESeq2, following the same procedures described above.

### Bulk RNA-seq data and epigenomic data processing

Raw RNA-seq data for AN and H9 naïve and primed human embryonic stem cells (hESCs) were retrieved from GEO (GSE138762 and GSE138688), including six naïve and two primed bulk RNA-seq libraries. Corresponding ATAC-seq bigWig files were downloaded from GSE138761. Transcript quantification and TSR usage analysis were performed using TSRdetector with the CHM13 genome assembly and RefSeq annotations. A TSR usage difference threshold of 25% was applied to identify regions with significant differential usage between naïve and primed hTSCs.

## ACKNOWLEDGEMENTS

This work was supported by the NIH Maximizing Investigators’ Research Award (R35 GM142917) and Chan Zuckerberg Initiative.

## AUTHOR CONTRIBUTIONS STATEMENT

S.F. and Z.B. conceived of and designed the study. S.F. and P.W. performed the data analysis. S.F. and B.Z. wrote the paper. B.Z. supervised the study.

## CODE AVAILABILITY

The available: github

